# Chemogenetic Inhibition of Infralimbic Prefrontal Cortex GABA-ergic Parvalbumin Interneurons Attenuates Chronic Stress Adaptions in Male Mice

**DOI:** 10.1101/792721

**Authors:** Nawshaba Nawreen, Rachel Morano, Parinaz Mahbod, Evelin M. Cotella, Khushali Dalal, Maureen Fitzgerald, Susan Martelle, Benjamin A. Packard, Rachel D. Moloney, James P. Herman

## Abstract

Hypofunction of the prefrontal cortex (PFC) contributes to stress-related neuropsychiatric illnesses. Mechanisms leading to prefrontal hypoactivity remain to be determined. Prior evidence suggests that enhanced activity of parvalbumin (PV) expressing GABAergic interneurons (INs) play a role in chronic stress related pathologies. In this study, the role of PFC PV INs in stress related phenotypes were explored using Cre inducible inhibitory DREADDs (Designer Receptors Exclusively Activated by Designer Drugs). Mice were first tested in the tail suspension test (TST) to determine the effects of PV IN inhibition during acute stress. Following this, the long term impact of PV IN inhibition during chronic variable stress (CVS) was tested in the forced swim test (FST). Acute PV IN inhibition reduced active (struggling) and increased passive coping behaviors (immobility) in the TST. In contrast, inhibition of PV INs during CVS increased active and reduced passive coping behaviors in the FST. Moreover, chronic inhibition of PV INs attenuated CVS-induced changes in Fos expression in the prelimbic cortex, basolateral amygdala and ventrolateral periaqueductal gray and also prevented adrenal hypertrophy and body weight loss associated with chronic stress. Our results suggest differential roles of PV INs to acute vs chronic stress, indicative of distinct biological mechanisms underlying acute vs. chronic stress responses. Our results also indicate a role for PV INs in driving chronic stress adaptation and support literature evidence suggesting cortical GABAergic interneurons as a therapeutic target in stress related diseases.

**SIGNIFICANCE:** Stress related diseases are associated with prefrontal hypoactivity, the mechanism of which is currently not known. In this study we showed that by inhibiting prefrontal GABA-ergic Parvalbumin interneurons (PV INs) using DREADDs, we can attenuate some of chronic stress related phenotypes. Additionally, we showed that modulation of PV IN activity during acute vs chronic stress had opposing effects on stress coping strategies, suggesting different underlying mechanisms behind acute vs chronic stress paradigms. Our findings indicate that GABA-ergic PV INs may be involved in driving stress related phenotypes and thereby an important target for treatment of stress-related illnesses. Our data suggest that reducing PV IN activity to promote prefrontal output may be an effective treatment strategy for stress related disorders.

## INTRODUCTION

Mood disorders (e.g., post-traumatic stress disorder (PTSD) and major depressive disorder (MDD) are associated with alterations in ventromedial prefrontal cortex (Broadman area 25) structure, activity and connectivity (Drevets et al., 2008; Hasler et al., 2008; Holmes et al., 2018; Murray et al., 2011; Rogers et al., 2004). To date, no universally efficacious therapeutic strategy exists for these neuropsychiatric conditions, despite having a lifetime prevalence of over 20% (Duman and Duman, 2015; Kessler et al., 2005). Studies in both humans and animal models have shown that chronic stress impairs functioning of the prefrontal cortex (PFC), potentially making prefrontal hypofunction a factor in the etiology of mood disorders (Drevets et al., 1997; Duman and Duman, 2015; Li et al., 2011; McKlveen et al., 2016; Radley et al., 2006).

Various clinical and preclinical studies implicate altered GABAergic circuitry and prefrontal hypofunction in the generation of depression in humans as well as depression-related behaviors in rodent chronic stress models (Duman, 2014; Luscher et al., 2011; McKlveen et al., 2016; Musazzi et al., 2015; Veeraiah et al., 2014). Recent functional and electrophysiological studies indicate increased infralimbic (IL) PFC (rodent homolog of the human ventromedial prefrontal cortex) GABAergic transmission (e.g., increased inhibitory synaptic drive and increased expression of GABA-ergic marker) following chronic stress, suggesting that enhanced interneuron (IN) activity may be involved in disruption of prefrontal cortical signaling (McKlveen et al., 2016; Shepard et al., 2016).

GABA-ergic parvalbumin interneurons (PV INs) synapse onto cell bodies of PFC pyramidal neurons and exert strong control over medial PFC (mPFC) output, maintaining appropriate excitatory/inhibitory (E/I) balance (Cardin et al., 2009; Courtin et al., 2014; Tremblay et al., 2016) and coordinating oscillatory activity (gamma oscillation) required for efficient PFC signaling. Consequently PV INs are well-positioned to play an important role in stress mediated prefrontal dysfunction (DeFelipe et al., 2013; Ferguson and Gao, 2018; Rymar and Sadikot, 2007; Sherwood et al., 2007). Increases in expression of PV and enhancement of glutamatergic transmission onto PV INs following chronic stress, are associated with prefrontal hypofunction, anxiogenesis and impaired coping behaviors in FST (Page et al., 2018; Shepard et al., 2016; Shepard and Coutellier, 2017). Reduction in PV expression in PFC is also associated with antidepressant efficacy (Ohira et al., 2013; Page and Coutellier, 2019; Zhou et al., 2015). In contrast to chronic, acute inhibition of PV INs in the PFC has opposing effects, resulting in increase in passive coping behavior such as learned helplessness in mice (Perova et al., 2015). This suggests differential role of PV INs in response to acute vs chronic stress, indicative of distinct brain circuitry being involved in modulating acute vs chronic stress-mediated phenotypes.

This study was designed to specifically investigate the role of IL PV INs in driving physiological and behavioral manifestations of acute and chronic stress-related phenotypes. We employed a chemogenetic strategy using DREADDs (Designer receptors Exclusively Activated by Designer Drugs) to specifically inhibit PV INs in the IL mPFC during exposure to acute stress and throughout exposure to chronic variable stress (CVS). Our results indicate that acute inhibition of PV INs increases passive and decreases active coping behavior in tail suspension test (TST). In contrast chronic inhibition of IL PV INs reduces passive and increases active coping behavior in the forced swim test (FST). PV IN inhibition blocks CVS mediated reduction in Fos expression in stress related brain regions downstream of the IL. Additionally we show that inhibition of PV INs prevents CVS induced physiological effects such as adrenal hypertrophy and body weight loss. These data suggest that IL PV INs play a role in driving behavioral, neuronal and physiological adaptations associated with chronic stress and also indicate differential role of these INs in the context of acute vs chronic stress related phenotypes.

## MATERIALS AND METHODS

### Mice

Male breeders from BL6 PV-Cre knock-in homozygous mice line (*Pvalb^tm1(cre)Arbr^*, JAX stock# 017320, Jackson Laboratories Maine, USA) were bred with WT C57BL/6J females (JAX stock# 000664) to generate an in-house colony of heterozygous PV-Cre C57BL/6J at the [Author University] animal housing facility. Mice were maintained under standard conditions (12/12h light/dark cycle, 22 +/− 1°C, food and water *ad libitum*; 4 mice per cage) in accordance with the [Author University] Institutional Animal Care and Use Committee, which specifically approved all acute and chronic stress regimens employed in this proposal. All protocols conformed to the Society’s Policies on the Use of Animals in Neuroscience Research. All experiments were performed on adult male mice (∼7.5 months of age at surgery).

### Stereotaxic Viral Vector Injection with AAV vectors

PV-Cre mice were anaesthetized with isoflourane, scalp shaved and placed in the stereotax frame. The incision site was disinfected using chlorohexidine and 70% ethanol. An incision at the midline was made using a single-edged blade. Cre-dependent adeno-associated virus 2 (AAV2) vectors AAV2-hsyn-DIO-hM4D(Gi)-mCherry (Gift from Bryan Roth; Addgene viral prep # 44362-AAV2) and AAV2-hsyn-DIO-mcherry (Gift from Bryan Roth; Addgene viral prep # 50459-AAV2) were injected bilaterally at a volume of 300nL (∼10^12^ genome copies/ml) into the IL mPFC. The coordinates used were as follows: (anterior/posterior range defined as +1.8 mm anterior to bregma, medial–lateral range defined as +/− 0.2 mm lateral to the midsagittal suture; dorsal–ventral range defined as −2.9 mm ventral to skull (Paxinos and Watson, 2007). Viruses were infused using a 2ul hamilton syringe at a rate of 60nl/min for 5 minutes. Following infusion, the injector was kept at the site for 8 minutes to allow for the virus to diffuse. The injection site was covered with gel foam and incision site sutured. 2.5mg/kg Meloxicam was administered for 3 days following surgery. Behavioral studies and stress protocols were initiated 3 weeks post injection to allow sufficient time for viral expression. Diagrammatic representation of experimental timeline is outlined in Figure.1

**Figure 1.**
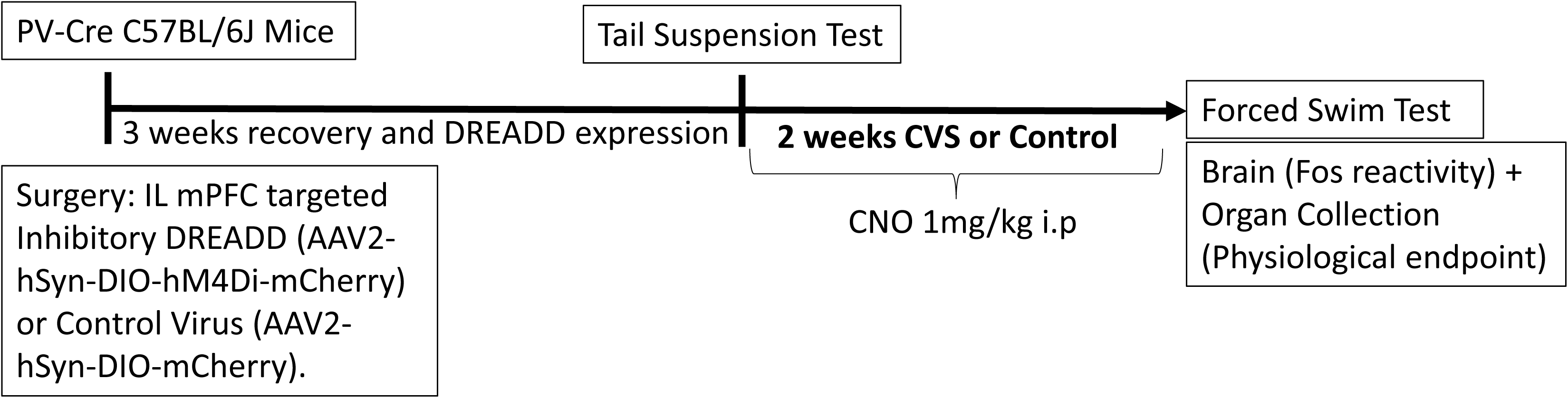
Experimental Design and Timeline. C57BL/6J PV-Cre mice approximately 7.5 months of age underwent surgery and were allowed 3 weeks to recover to enable sufficient time for DREADD expression. Animals were then subjected to CVS procedure twice a day for 14 days or served as controls. The first stressor was a tail suspension test (TST) to determine acute effects of PV IN inhibition in animals within the CVS group. Animals were dosed with 1mg/kg Clozapine N-oxide (CNO) prior to each stressor to inhibit PV INs during the CVS procedure. 24 hours after end of CVS, animals were subjected to forced swim test (FST) following which mice were euthanized, body was perfused, and brains and organs were collected. No CNO was administered during the testing phase in FST.

### Chronic Variable Stress (CVS) Procedure

During the CVS procedure, mice were subjected to a series of randomly alternating stressors administered twice daily over a period of 14 days. The unpredictable stressors used were as follows: restraint (30 minutes), cold room exposure (15 min, 4°C), shaker stress (1 hour, 100 rpm), hypoxia (30 minutes, 8% oxygen and 92% nitrogen), Y maze (8 minutes), shallow water (30 min), wet bedding (2 hours) and cage tilt (2 hours, 45°). Body weights were measured on days 1, 4, 8 and 14 during the CVS procedure.

### Drug Administration

Clozapine N-oxide (CNO, NIMH Chemical synthesis and Drug Supply Program) was used as the DREADD actuator to activate the inhibitory DREADD. CNO was dissolved in 5% dimethyl sulfoxide (Sigma) and then diluted with 0.9% saline and administered intraperitoneally at a dose of 1mg/kg twice a day, 30 minutes before start of each stressor. All animals received chronic injection of CNO for 14 days. A maximum time period of 6 hours was given between stressors, to allow sufficient time for CNO to be cleared from the body (Jendryka et al., 2019; MacLaren et al., 2016). Each individual stress session did not last for more than 2 hours in order to ensure that CNO was on-board throughout the stressor.

### Behavioral Assessments

#### Tail Suspension Test (TST)

The TST (Can et al., 2012) was used as the first stressor in the CVS group to observe acute effects of inhibiting PV INs on passive coping behavior. Mice were suspended 55 cm above ground using a 17cm long tape that was attached to a suspension bar, for a total time period of 6 minutes. Sessions were video-recorded from the side to allow full body visualization of mice behaviors-active coping (struggling) behavior, which comprised of strong shaking of the body and movement of all 4 limbs, and passive coping (immobility) behavior which comprised of not making any active limb movements. Latency to reach immobility was also measured. Behaviors were quantified by an experimenter blinded to the group assignments using behavioral scoring software Kinoscope 3.0.4. Behaviors during the 6 minute block were reported.

#### Forced Swim Test (FST)

FST was conducted as a novel stressor 24 hr following completion of the CVS procedure. Mice were placed in a clear cylinder ( 2L glass beaker) filled with water (24±1°C, 18 cm depth) for a period of 10 minutes. Sessions were video-recorded from the side to allow full body visualization for total immobility duration, which comprised of not making any active movements or floating in the water without struggling, and total swimming duration, which comprised of moving limbs in an active manner and making circular movements around the cylinder. Behaviors were quantified by an experimenter blinded to the group assignments using behavioral scoring software Kinoscope 3.0.4. Behaviors during the 10 minute block were reported. To control for a potential effect of chronic exposure to CNO on FST, a separate group of PV-Cre mice (n=4) was chronically injected with saline for 14-days (no CVS) and then behaviorally tested in the FST in parallel with the rest of the animals.

### Euthanasia and Tissue Collection

Mice were euthanized with an overdose of sodium pentobarbital after FST, and transcardially perfused with 0.9% saline followed 4% paraformaldehyde in 0.01 M phosphate buffer (PBS), pH 7.4. Brains were removed and post-fixed in 4% paraformaldehyde at 4°C for 24 hours, then transferred to 30% sucrose in 0.01MPBS at 4°C until processed. Thymi and adrenal glands were collected, cleaned and weighed from all animals.

### Immunohistochemistry

Brains were sectioned into 30μm coronal sections using a freezing microtome (−20°C). Sections were collected into 12 wells (1/12) containing cryoprotectant solution (30% Sucrose, 1% Polyvinyl-pyrolidone (PVP-40), and 30% Ethylene glycol, in 0.01 M PBS). Immunohistochemistry was performed at room temperature and 0.01M PBS was used to rinse brain slices before each treatment described below.

#### Injection site and viral spread

To determine if virus spread was restricted to the IL, sections were incubated with a rabbit anti-mCherry (1:500, Abcam, ab167453) for 2 hours, followed by visualization with donkey anti-rabbit Cy3 conjugate (1:500, Invitrogen, A10520). Images were acquired using Nikon Confocal Microscope at 2.5X and 5X magnification.

#### PV and DREADD Co-localization

Targeting of IL mPFC PV neurons and recombination of hM4Di DREADD was verified by co-localization of PV immunoreactivity with virally expressed mCherry fluorescence. Free floating sections were incubated in blocking solution (4% normal goat serum (NGS), 0.1% TritonX-100, 0.1% bovine serum albumin (BSA) in 0.01M PBS) for 1 hour at RT. After that, sections were incubated with rabbit anti-PV (1:1000, Abcam, ab11427) overnight, followed by visualization with donkey-anti-rabbit Alexa 488 conjugate (1:500, Invitrogen, A11034). Images were acquired using Nikon Confocal Microscope at 40X magnification.

#### Fos Immunoreactivity

Neuronal activation was measured using Fos as a marker. Free floating sections were incubated in 1% Sodium Borohydride for 20 minutes and then in 3% hydrogen peroxide in PBS for 20 minutes. After that, slices were incubated in blocking solution (NGS, 0.3% TritonX-100, 0.2% bovine serum albumin (BSA) in 0.01M PBS) for 1 hour. Sections were then incubated with Fos rabbit polyclonal antibody (1:200, Santa Cruz, sc-52) in blocking solution overnight and was followed by incubation in secondary antibody (biotinylated goat anti-rabbit, (1:400; Vector Laboratories, BA1000) in blocking solution for 1 hour the next day. Sections were then treated with avidin-biotin horseradish peroxidase complex (1:800 in 0.01M PBS; Vector Laboratories, PK6100) for 1 hour and then developed with an 8 minute incubation in DAB-Nickel solution: 10mg 3,3′-diaminobenzidine (DAB) tablet (Sigma, DF905), 0.5 ml of a 2% aqueous nickel sulfate solution, 20ul of 30% hydrogen peroxide in 50ml of 0.01M PBS. Sections were mounted on superfrost slides (Fisherbrand, Fisher), allowed to dry, dehydrated with xylene, and then coverslipped with DPX mounting medium (Sigma).

Images were acquired using microscope Carl Zeiss Imager Z1 at a 5X objective. For analysis, we counted minimum of 3 bilateral sections per brain region/animal covering the prelimbic cortex (PrL) (Bregma 2.80mm to 1.98mm), basolateral amygdala (BLA) (Bregma −1.06 to −1.58) and ventrolateral periaqueductal gray (vlPAG) (Bregma −4.16 to −4.36) as defined in the Franklin and Paxinos mouse brain atlas (3^rd^ Edition) (Franklin and Paxinos, 2008). The number of Fos positive nuclei was counted using a semi-automated analysis macro in the Image J software package (National Institutes of Health, Bethesda, MD). The macro was generated using the “Analyze Particle” tool, with a defined common level of background intensity, nuclei circularity and size (previously validated manually). The relative density of the population of immunopositive cells was calculated by dividing the number of Fos positive cells by the respective brain area.

### Statistical Analysis

The experiment was setup as a 2X2 study design, with stress (CVS or No CVS) and DREADD (hM4Di or control) as factors with a sample size of n=10 per group (See Figure 1 for experimental design and timeline). Statistical analyses for FST and Fos protein quantification were performed using a two-way ANOVA with stress (No CVS, CVS) and DREADD (Control, hM4Di) as main factors. TST data were analyzed using Student’s t-test. FST measurements over time were done using two-way repeated measure ANOVA with stress (No CVS, CVS) and DREADD (Control, hM4Di) as main factors analyzed over time. Tukey’s post-hoc test was performed in cases with significant interaction between factors. Because specific hypotheses were formed *a priori* on the effects of CVS within groups, planned comparisons using Fisher’s least significant difference (LSD) were performed in cases with no significant interaction effect. Data were analyzed by STATISTICA 7.0 (Statsoft, Inc.,Tulsa, USA) and Graph Pad Prism 8.1.2 (GraphPad Software, La Jolla California USA). Outliers were detected using the Grubbs’ test (GraphPad Software) and removed from analysis. After exclusion of outliers, data were assessed for normal distribution (Shapiro-Wilk) and appropriate parametric and /or non-parametric tests used. Data are presented as mean ± SEM with statistical significance set at *P* ≤ 0.05. See Table 1 for details regarding data structure and type of test used. Superscript letters listed with p-values correspond to the statistical tests shown in Table 1.

**Table 1.**
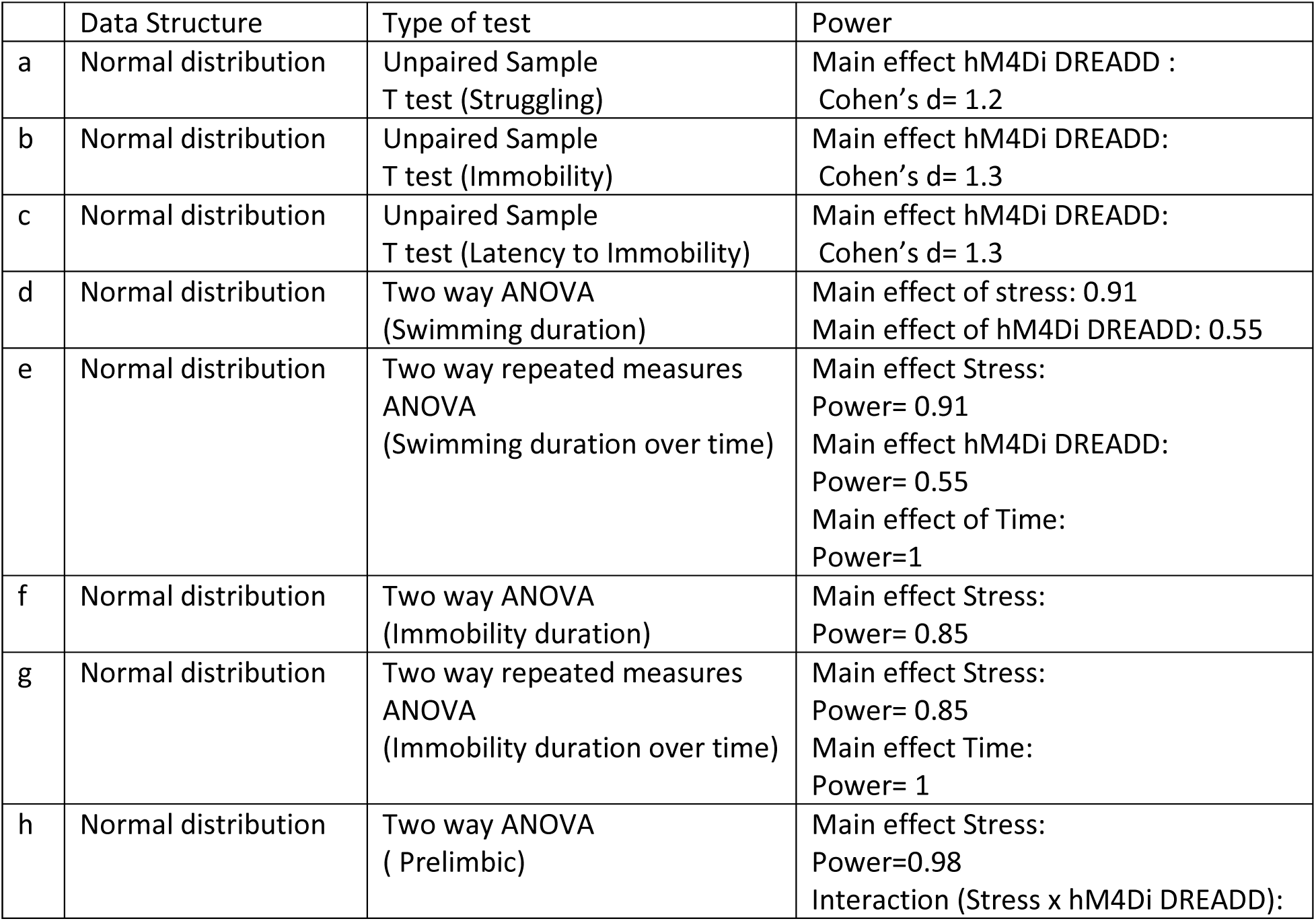
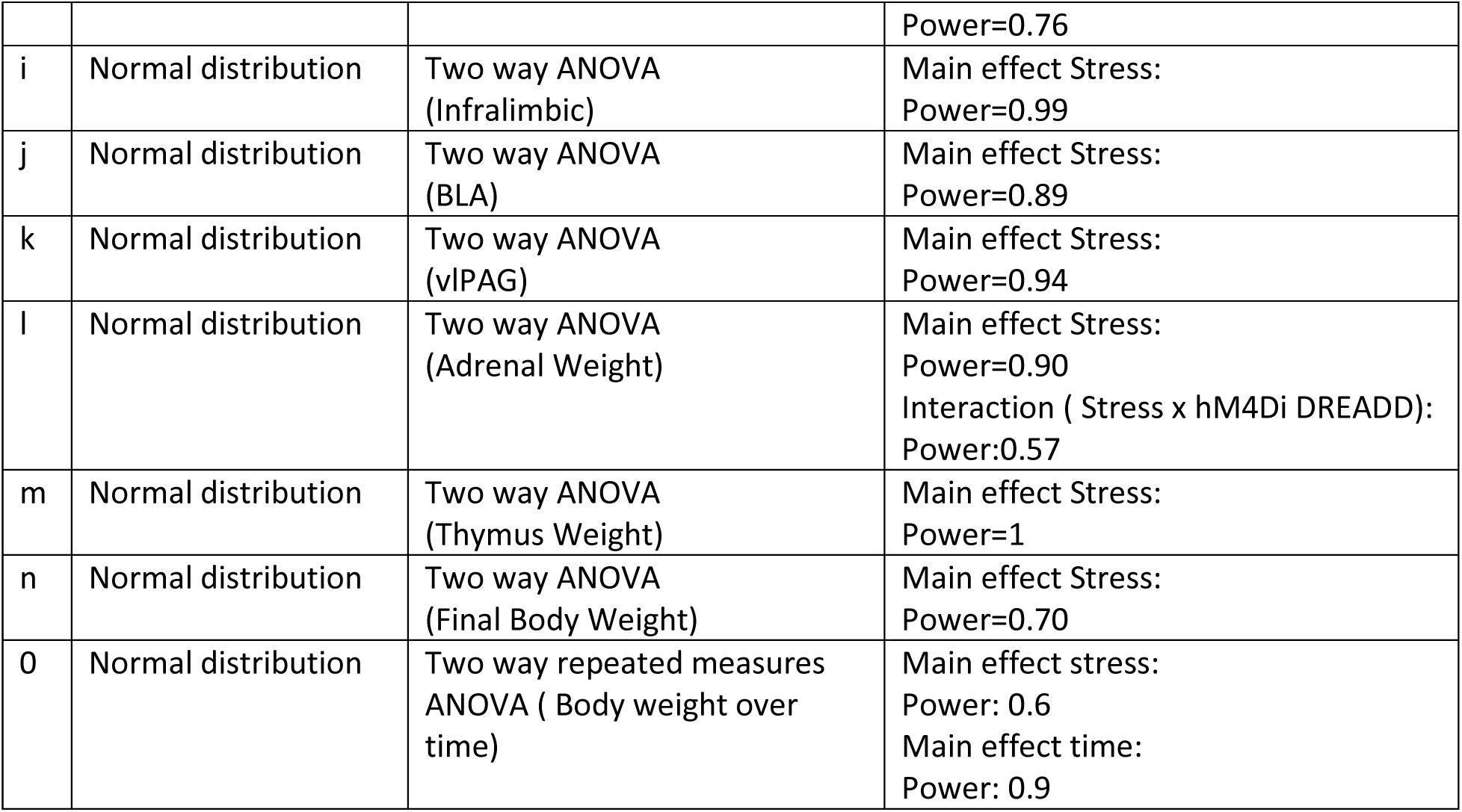
Data Structure, type of test to analyze the data, and observed power of key results.

|  | Data Structure | Type of test | Power |
| --- | --- | --- | --- |
| a | Normal distribution | Unpaired Sample T test (Struggling) | Main effect hM4Di DREADD :<br>Cohen's d= 1.2 |
| b | Normal distribution | Unpaired Sample T test (Immobility) | Main effect hM4Di DREADD:<br>Cohen's d= 1.3 |
| c | Normal distribution | Unpaired Sample T test (Latency to Immobility) | Main effect hM4Di DREADD:<br>Cohen's d= 1.3 |
| d | Normal distribution | Two way ANOVA (Swimming duration) | Main effect of stress: 0.91<br>Main effect of hM4Di DREADD: 0.55 |
| e | Normal distribution | Two way repeated measures ANOVA (Swimming duration over time) | Main effect Stress:<br>Power= 0.91<br>Main effect hM4Di DREADD:<br>Power= 0.55<br>Main effect of Time:<br>Power=1 |
| f | Normal distribution | Two way ANOVA (Immobility duration) | Main effect Stress:<br>Power= 0.85 |
| g | Normal distribution | Two way repeated measures ANOVA (Immobility duration over time) | Main effect Stress:<br>Power= 0.85<br>Main effect Time:<br>Power= 1 |
| h | Normal distribution | Two way ANOVA ( Prelimbic) | Main effect Stress:<br>Power=0.98<br>Interaction (Stress x hM4Di DREADD): |
|  |  |  | Power=0.76 |
| i | Normal distribution | Two way ANOVA (Infralimbic) | Main effect Stress:<br>Power=0.99 |
| j | Normal distribution | Two way ANOVA (BLA) | Main effect Stress:<br>Power=0.89 |
| k | Normal distribution | Two way ANOVA (vIPAG) | Main effect Stress:<br>Power=0.94 |
| l | Normal distribution | Two way ANOVA (Adrenal Weight) | Main effect Stress:<br>Power=0.90<br>Interaction ( Stress x hM4Di DREADD):<br>Power:0.57 |
| m | Normal distribution | Two way ANOVA (Thymus Weight) | Main effect Stress:<br>Power=1 |
| n | Normal distribution | Two way ANOVA (Final Body Weight) | Main effect Stress:<br>Power=0.70 |
| o | Normal distribution | Two way repeated measures ANOVA ( Body weight over time) | Main effect stress:<br>Power: 0.6<br>Main effect time:<br>Power: 0.9 |

## RESULTS

### Selective Targeting of PV INs Achieved Using DREADDs

The AAV constructs used in this experiment are shown in Figure 2A. Following Cre recombination, the viral construct expresses inhibitory DREADD sequence hM4Di along with a fluorescent reporter (mCherry), allowing visualization of cells undergoing recombination. Control virus was a Cre-inducible mCherry lacking the DREADD hM4Di construct. Figure 2B depicts stereotaxic injection site in the IL and also demonstrates that the expression of the DREADD was restricted to the IL. Cre mediated recombination of hM4Di-mCherry and cell type specificity were conferred by immunostaining. Expression of red hM4Di-mCherry demonstrates successful recombination (Figure 2C-left). hM4Di-mCherry expression was restricted to PV INs only, confirmed by colocalization of red mCherry with green PV immunostaining resulting in yellow overlay (Figure 2C-right).

**Figure 2.**
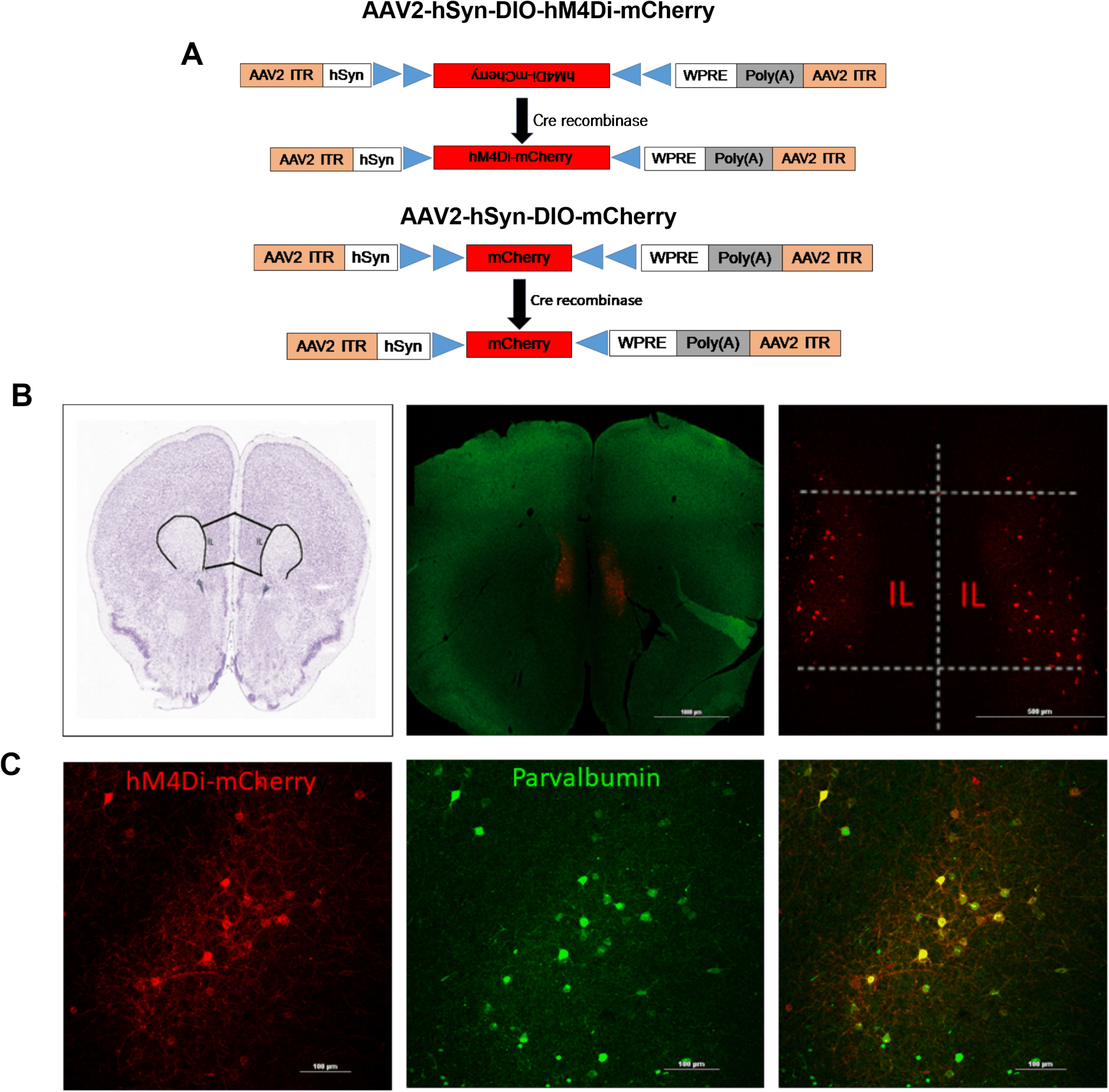
Targeting PV INs in the mPFC using DREADDs. Panel A represents the design of AAV2-hSyn-hM4Di-mCherry (top) and AAV2-hSyn-mCherry (bottom) vectors employing the DIO strategy. Two pairs of heterotypic, antiparallel loxP recombination sites (blue triangles) achieve Cre-mediated transgenes inversion and expression under the control of hSyn promoter. ITR: left-inverted terminal repeat, hSyn: human synapsin, WPRE: woodchuck hepatitis DREADD post-transcriptional regulatory element. Panel B represents schematic coronal section illustrating the injection site of the imaged area in the PFC. mCherry fluorescence was detected in the IL following bilateral injection into PV-Cre mice. Scale bar: 1000 and 500 μm. Panel C represents successful Cre mediated recombination of DREADDs demonstrated by presence of red mCherry (left); green fluorescence identifies PV INs (middle); hM4Di receptors selectively expressed in PV INs as illustrated by mCherry (red) and PV (green) co-expression (overlay, yellow, right image) Scale bar: 100 μm.

### Acute Inhibition of PV INs: TST

The behavioral consequences of acute inhibition of PV INs in the IL were tested by using the TST as the first stressor in the CVS paradigm (Figure 3). Animals were dosed with 1mg/kg CNO 30 minutes prior to the start of TST. We observed significant changes in coping behavior following acute inhibition of PV INs in the IL. Compared with control mice, mice expressing hM4Di showed significant reduction in struggling duration (t= 2.7, df=18, p=0.02^a^; Figure 3A), significant increase in immobility duration (t=2.9, df=18, p=0.009^b^; Figure. 3B) and decreased latency to immobility (t=2.5, df=17, p=0.02^c^; Figure. 3C) respectively.

**Figure 3.**
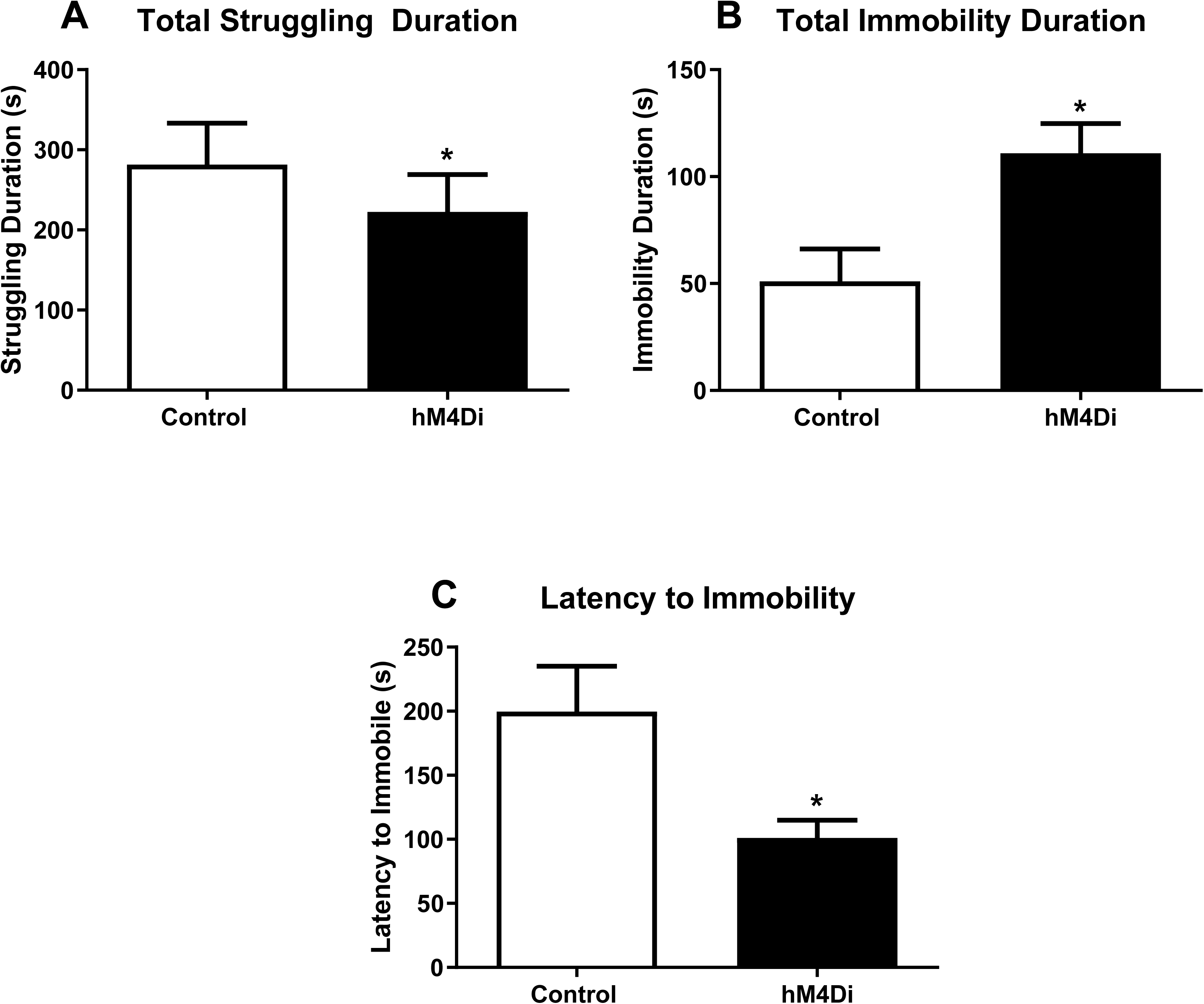
Impact of acute chemogenetic inhibition of PV IN in the IL mPFC on coping behavior in the tail suspension test (TST). Acute inhibition of PV INs in the IL mPFC during TST, reduced total time spent struggling (A), increased total time spent immobile (B) and reduced latency to immobility (C). All mice were treated with CNO [1mg/kg, intra peritoneal (i.p.)] 30 minutes before TST. Behaviors were analyzed for a total time of 6 minutes. Values represent mean ± SEM, n = 9–10 per group. * indicates significant effect p < 0.05 versus corresponding control group.

### Chronic Inhibition of PV INs during CVS: Impact on FST

Animals were tested for coping behaviors in the FST the day following cessation of CVS. Our purpose was to test whether inhibition of PV INs during the chronic stress regimen, could block the aggregate effect of repeated stress on subsequent coping behavior (FST used as a novel stressor) and brain activation patterns (Fos expression). All subjects received viral (Control or hM4Di) and CNO treatments, and because CNO was only administered during CVS, any phenotypes observed during FST were interpreted as reflecting an impact of PV IN manipulation during CVS on subsequent stress coping behavior. We observed significant differences in FST coping behaviors following chronic inhibition of PV INs during CVS. Specifically, chronic PV IN inhibition during stress increased active coping (swimming) and reduced passive coping (immobility) behaviors in the FST. Two-way ANOVA of total swimming duration showed a significant main effect of stress [F(1,35)=11.7; p=0.002^d^; Figure 4A] and DREADD [F(1,35)=4.7; p=0.037^d^)] but no stress x DREADD interaction [F(1,35) =1.47; p=0.2]. Planned comparisons revealed a significant increase in swimming duration in the CVS hM4Di group compared with both CVS Control (p=0.02; Figure 4A) and No CVS hM4Di group (p=0.002; Figure 4A). Analysis of swimming behavior over time showed a main effect of stress [F(1,35)=11.7; p=0.002^e^; Figure 4C], DREADD [F(1,35)=4.7; p=0.037^e^)] and time [F(9,315)=68.2; p<0.0001^e^)] but no interaction effects were observed among the 3 groups time x stress x DREADD [F(9,315)=0.6; p=0.78]. Planned comparisons revealed significant increase in swimming duration in the CVS hM4Di group at 2, 3, 4 and 8 minutes timepoints compared with No CVS hM4Di group (p=0.004,0.009,0.01 and 0.009 respectively; Figure 4C).

**Figure 4.**
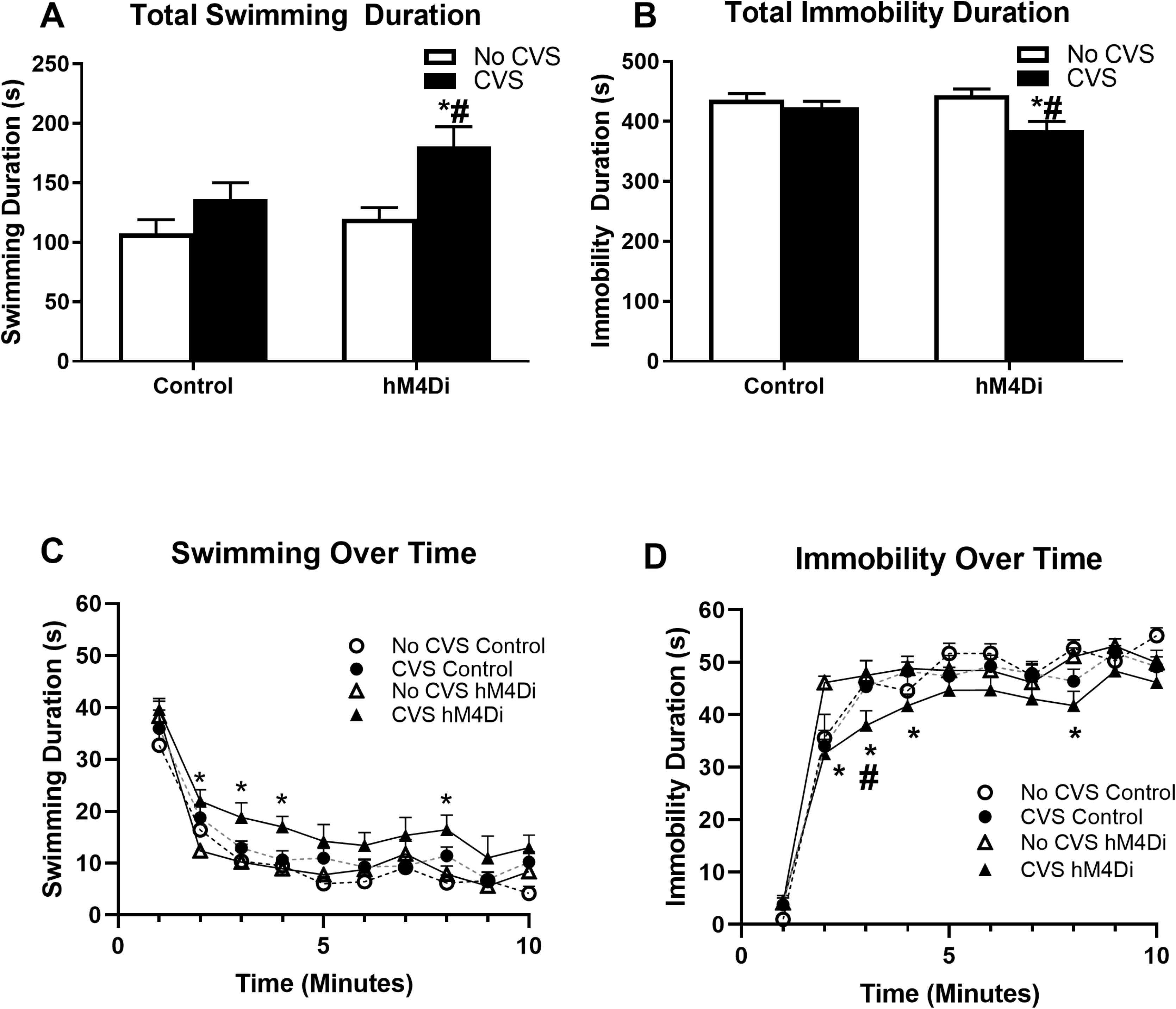
Effects of chronic chemogenetic inhibition of PV INs during CVS on coping behavior in the forced swim test (FST) following CVS. Chronic inhibition of PV INs in the IL mPFC during CVS, resulted in increased total time spent swimming (A) and decreased total time spent immobile (B) in the FST. C and D represent changes in swimming and immobility behavior respectively over the 10 minutes of FST. Values represent mean ± SEM, n = 9–10 per group. * indicate planned comparison significant effect p<0.05 versus corresponding No CVS hM4Di group. # indicates planned comparison significant effect p<0.05 versus corresponding CVS Control group.

Two-way ANOVA of total immobility duration showed a significant main effect of stress [F(1,35) =9.5; p=0.004^f^; Figure 4B], no main effect of DREADD [F(1,35)=1.7; p=0.2)] and no stress x DREADD interaction [F(1,35)=3.9; p=0.057]. Planned comparisons revealed a significant reduction in immobility duration in the CVS hM4Di group compared with CVS Control (p=0.02) and No CVS hM4Di group (p=0.0009). There was a significant main effect of stress [F(1,35)=9.5; p=0.004^g^; Figure 4D)] and time [F(9,315)=156.4; p<0.0001^g^)] on immobility duration but no interaction effects were observed among the 3 groups time x stress x DREADD [F(9,315)=1.1; p=0.4]). Planned comparisons revealed a significant decrease in immobility in the CVS hM4Di group at 2, 3, 4 and 8 minute timepoints compared with No CVS hM4Di group ( p=0.00009;0.005;0.04;0.007 respectively; Figure 4D) and at the 3 minute timepoint compared with the CVS Control group ( p=0.03; Figure.4D). Chronic PV IN inhibition had no effect on locomotor activity, demonstrated by no significant difference in total distance travelled (t=0.74; df=18; p=0.46; Figure 4-1A) or velocity (t=0.75; df=18; p=0.47; Figure 4-1B) demonstrating behavioral effects were not confounded by locomotor deficits. Control experiments to determine effects of chronic CNO in FST behavior, showed no significant difference in immobility duration in CNO vs saline group (t=1.39; df=10; p=0.2; Figure 4-2).

### Chronic Inhibition of PV INs during CVS: Impact on Fos Induction by FST

To test for Fos activation, animals were perfused after FST and brains were collected to analyze neuronal activation in brain regions typically activated by stress. We observed significant reduction in Fos induction in the CVS Control group compared to No CVS Control group, in the PrL, IL, BLA and vlPAG. Inhibition of PV INs during CVS prevented the reduction in Fos expression caused by CVS in the PrL, BLA and vlPAG, but not in the IL. Analysis of the PrL revealed a significant main effect of stress [F(1,30)= 17.5; p=0.0002^h^; Figure 5A] and a significant stress x DREADD interaction [F(1,30)= 7.7; p=0.009^h^]. Post hoc analysis revealed a significant reduction in Fos expression in the CVS Control group (p=0.0003) which was prevented by chronic PV IN inhibition in the CVS hM4Di group (p=0.75). There was a significant main effect of stress only [F(1,27)= 21.2; p<0.0001^i^, Figure 5B] on Fos expression in the IL. There was a significant main effect of stress in the BLA [F(1,12)= 12.4; p=0.004^j^; Figure 5C] and vlPAG [F(1,12)= 16.4; p=0.004^k^; Figure 5D] as well, with planned comparisons revealing significant reduction in Fos expression in the CVS Control group (p=0.008 and p=0.005 in BLA and vlPAG respectively) that was prevented by chronic PV IN inhibition in the CVS hM4Di group (p=0.75 and p=0.6 in BLA and vlPAG respectively). Analysis of Fos protein expression in the lateral septum (LS), anterior and ventral bed nucleus of the stria terminalis (BNST) and dorsolateral PAG (dlPAG) showed no significant treatment effects of PV IN inhibition demonstrating those regions were not affected by PV IN modulation (Table 5-1).

**Figure 5.**
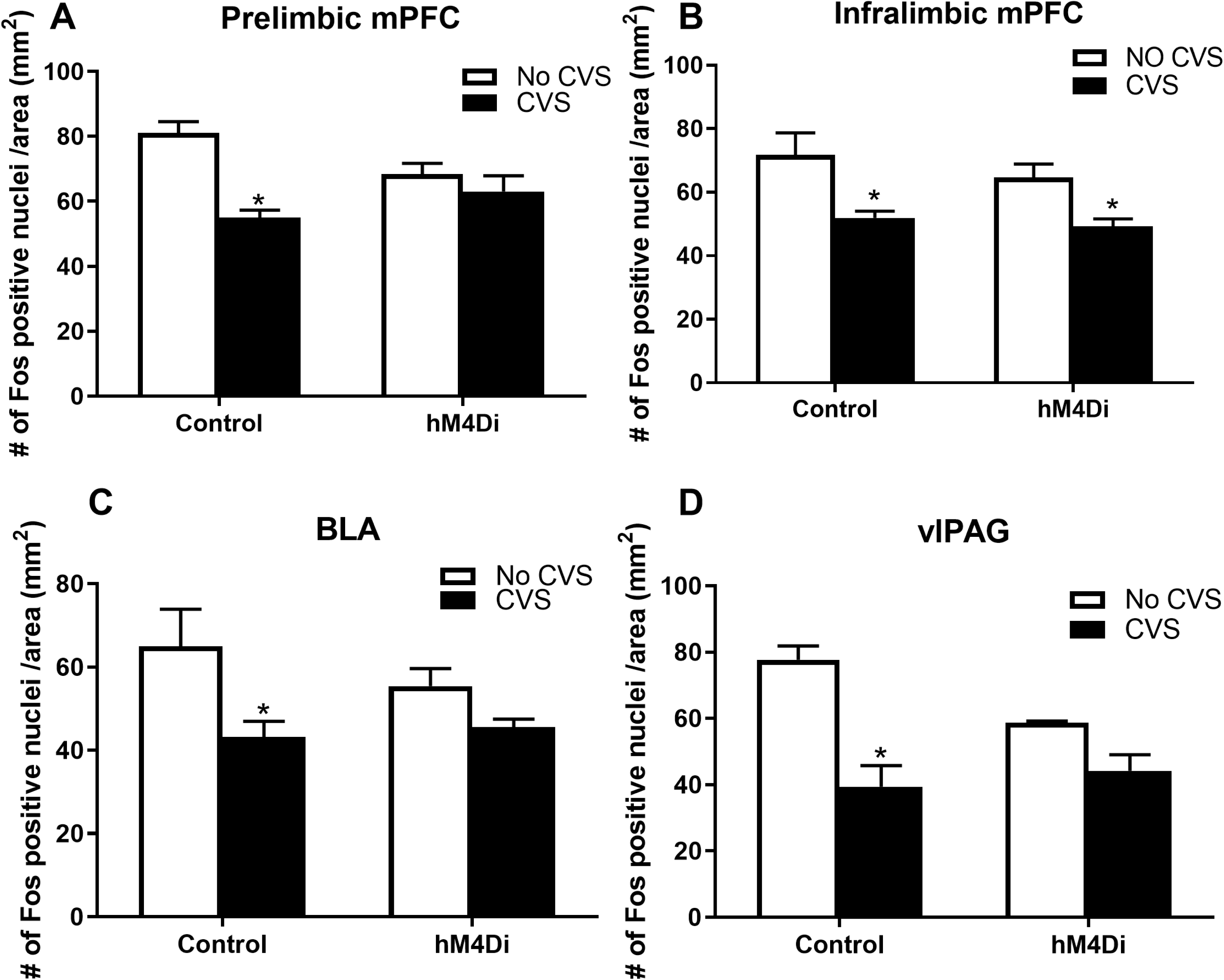
Fos immunoreactivity in the PrL, IL, BLA and vlPAG. Chronic inhibition of PV INs in the IL mPFC during CVS, prevented CVS mediated reduction in Fos expression in the Prelimbic cortex, BLA and vlPAG respectively (A, C and D) but did not prevent CVS mediated reduction in Fos expression in the infralimbic cortex (B) following FST. Data are presented as mean ± s.e.m. * indicates significant result p < 0.05 post hoc (A) and planned comparisons (C and D) compared to respective No CVS Control groups.

### Chronic Inhibition of PV INs during CVS: Impact on Physiological Outcomes

Organs and body weights were used to assess somatic effects of CVS. Adrenal hypertrophy and/or thymic atrophy are often observed following chronic stress and are used as indicators of repeated/chronic hypothalamic pituitary adrenal (HPA) axis activation. Here, there was a main effect of stress [F(1,35)=11.2; p=0.002^l^; Figure 6A] and a significant stress x DREADD interaction [F(1,35)=4.8; p=0.035^l^] on adrenal gland weights. Post hoc analysis using Tukey’s test revealed that CVS Control group had significantly increased adrenal weight compared to No CVS Control group (p=0.002), which was prevented by CVS hM4Di when compared to No CVS hM4Di group (p=0.84). There was a main effect of stress on thymus weight [F(1,35)=161.4; p<0.0001^m^; Figure 6B], with no effect of DREADD [F(1,35)=0.2; p=0.7] or stress X DREADD interaction [F(1,35)=57; p=0.1].

**Figure 6.**
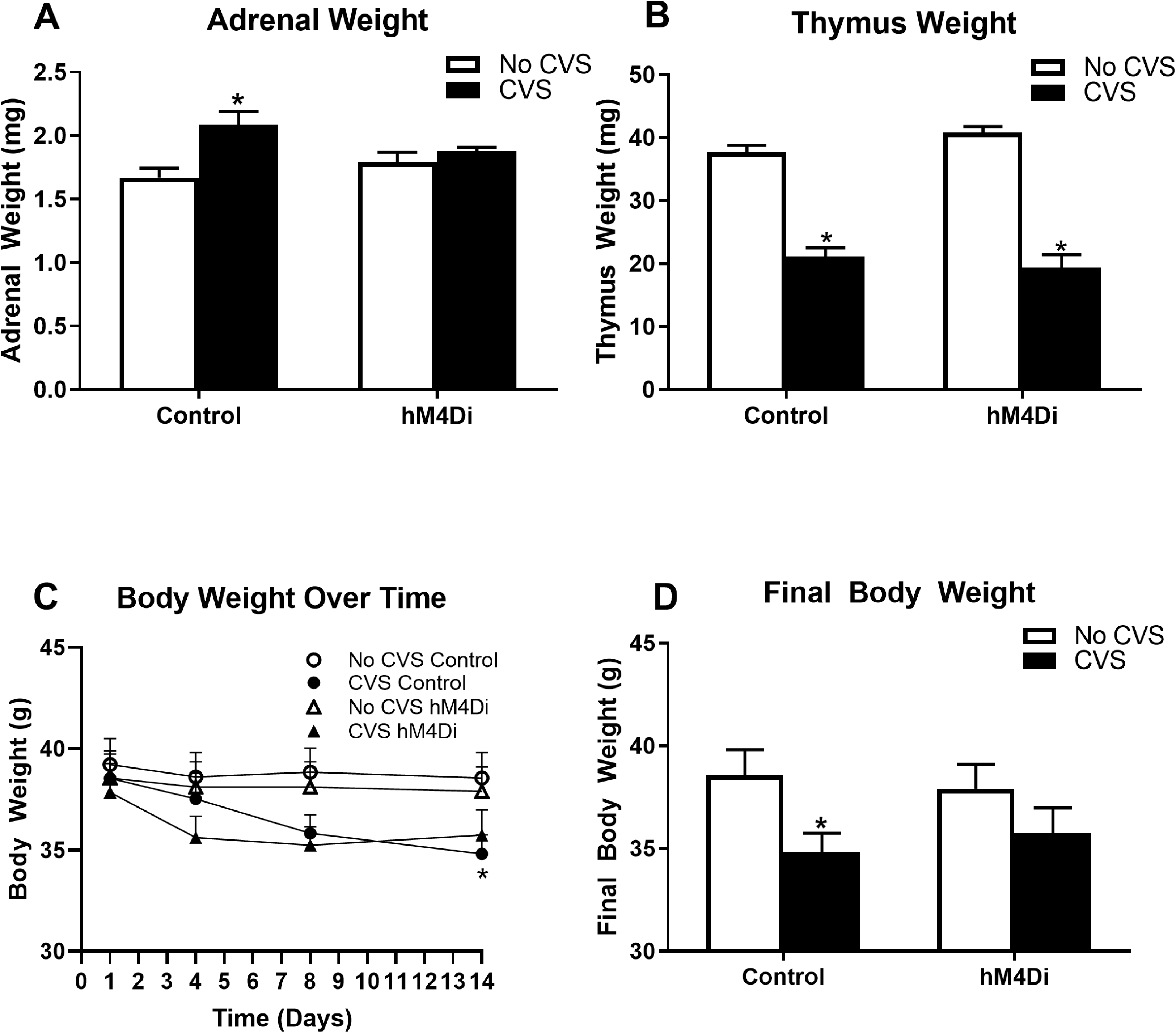
Impact of chronic stress on organ and body weights. Chronic stress resulted in increase in adrenal gland weight in CVS Control group only (A); decrease in thymus size in both CVS Control and hM4Di groups (B); decrease in body weight over time in CVS Control (C) and decrease in final body weight in CVS Control group only (D). Data are presented as absolute organ and body weights. Values represent mean ± SEM, n = 9–10 per group. * indicate planned comparisons significant effect p < 0.05 versus corresponding No CVS Control groups.

Decreased body weight gain is observed following chronic mild stress exposure in rodents (Ghosal et al., 2017). Two-way repeated measures ANOVA of body weight over time during the 14 days CVS paradigm showed a main effect of stress [F(1,36)= 5.3, p=0.03°] and time [F(3,108)=4.5; p=0.005°; Figure. 6C]. Planned comparisons revealed body weight in CVS Control group to be significantly lower than No CVS Control only on day 14 (p=0.03) (Figure.6C). We observed a significant main effect of stress on final body weight [F(1,35)=6.6; p=0.01^n^; Figure. 6D] with no effect of DREADD [F(1,35)=0.2; p=0.6] or stress x DREADD interaction [F(1,35)=0.006; p=0.9], consistent with known effects of CVS on body weight gain. Planned comparisons revealed final body weight in CVS group to be significantly lower than No CVS Control group (p=0.03), which was prevented by CVS hM4Di when compared to No CVS hM4Di group (p=0.08) (Figure.6D).

## DISCUSSION

Our studies support a role for prefrontal GABAergic PV INs in acute and chronic stress-mediated behavioral and physiological phenotypes. Acute inhibition of PV INs in stress naive animals resulted in an increase in passive coping and a decrease in active coping in the TST. In contrast, chronic inhibition of PV INs during CVS resulted in a decrease in passive coping and an increase in active coping strategy during FST, suggestive of dynamic behavioral remodeling during an aversive challenge. Chronic stress-induced behavioral alterations were accompanied by changes in neuronal activation patterns following FST, quantified by Fos expression in regions downstream of the IL. Chronic PV IN inhibition prevented CVS induced reductions in Fos expression in PrL, BLA and vlPAG, indicating that inhibition of PV INs mitigates the impact of chronic stress on stress regulatory brain regions. PV IN inhibition during CVS also prevented CVS induced adrenal hypertrophy and body weight loss, further suggesting PV IN signaling during CVS might be playing a role in physiological effects of chronic stress. Overall, the data indicate that PV INs may play a role in inhibiting IL output during chronic stress, suggesting a potential role in driving ventromedial PFC hypofunction.

GABAergic PV INs are well positioned to provide strong, fast-spiking inhibitory signals to pyramidal projection neurons in the PFC and reduce network excitability, and therefore could be contributing to chronic stress-mediated hypoactivity (Safari et al., 2017; Tremblay et al., 2016; Winkelmann et al., 2014). Our data suggest that PV INs play an important role in chronic stress-mediated inhibition of the IL. Chronic inhibition of IL PV INs during CVS resulted in increased active and decreased passive coping behaviors in FST. A switch to active coping can be interpreted as an adaptive strategy to deal with chronic stress, and drugs that are effective antidepressants in humans typically promote active coping styles and reduce passive coping in the FST in mice (Martí and Armario, 1993; Porsolt et al., 1977). GABA receptor antagonists have been shown to have antidepressant and anxiolytic properties (Bhutada et al., 2010; Mehta et al., 2017; SAMAD et al., 2018; Zanos et al., 2017). It is known that antidepressants such as fluoxetine and ketamine reduce PV expression in the PFC (Ohira et al., 2013; Page and Coutellier, 2019; Zhou et al., 2015). Moreover, preventing the reduction in PV IN activity leads to loss of antidepressant efficacy, further suggesting that reduced activity of PV INs might be playing a role in therapeutic efficacy of antidepressants (Page and Coutellier, 2019; Zhou et al., 2015). Therefore based on prior studies and our findings, inhibition of PV INs during chronic stress may lead to more adaptive stress coping strategies and reverse some of the behavioral deficits associated with chronic stress. It is important to note that the effects observed in FST are specifically due to changes in coping strategies due to PV IN inhibition and not due to any changes in locomotor activity (Figure 4-1). Additionally, we did not detect any effects on FST due to chronic dosing of CNO alone, suggesting that repeated CNO dosing did not alter stress coping behavior (Figure 4-2).

Our experiments revealed that inhibition of PV IN in the IL during stress can prevent chronic stress-induced decreases in Fos expression in key stress regulatory regions such as the PrL, BLA and vlPAG following FST (Berton et al., 2007; Keedwell et al., 2005; Vialou et al., 2014). Since we cannot verify direct PV IN modulation of IL projections to these regions, we cannot preclude the possibility that reversal of CVS-related inhibition of Fos induction is due to actions of the IL through other projection systems. Nonetheless, the data suggest that PV IN inhibition reduces inhibitory effects of CVS on IL outflow, permitting drive of downstream structures known to participate in physiological reactivity and stress coping behavior (Maier and Watkins, 2010). Notably, this includes the neighboring PrL, which is not targeted by our DREADD injections and thus has Fos excitability modulated by cortico-cortical connections. Involvement of the PrL is consistent with its prominent role in mediation of coping behavior (Fiore et al., 2015; Johnson et al., 2019; Molendijk and de Kloet, 2019).

Repeated inactivation of PV INs during stress prevented the CVS induced increase of adrenal weight. The adrenal glands are highly sensitive to repeated stress, and it is believed that increased adrenal size is linked to cumulative increases in ACTH secretion. Blockade of adrenal hypertrophy suggests that PV INs participate in control of the central limb of HPA axis activation and provides additional confirmation of cumulative efficacy of chronic PV IN inhibition in control of stress endpoints. In contrast to the adrenals, CVS caused equivalent decreases in thymus weight, suggesting either sensitization of glucocorticoid sensitivity or enhanced autonomic activation by CVS, presumably mediated by mechanisms independent of PV INs. Chronic inhibition of PV IN also prevented CVS induced reduction in final body weight, suggesting PV IN signaling might also be playing in a role in CVS mediated body weight effects.

As part of our design, we assessed the impact of IL PV IN inhibition acutely following the first stressor in our CVS regimen, the TST, which allows for a behavioral readout (duration of struggling, immobility and latency to immobility). Acute inhibition of IL PV INs resulted in decreased active coping (struggling) and increased passive coping (immobility) in the TST. Our data are consistent with a prior study indicating that reduced excitatory synaptic drive onto PV INs is linked with increase stress susceptibility and enhanced helplessness behavior (Perova et al., 2015). These data indicate that PV INs may play a role in driving active coping responses, when an animal with no history of prior stress is exposed to a novel acute stressor such as the TST. Together, these studies suggest that activation of PV INs is required for coping responses to acute stress.

Our results with acute PV IN inhibition are in contrast to the results seen in the FST after chronic PV IN inhibition during a two-week CVS exposure. These data indicate different roles for these neurons in acute vs. chronic stress adaptations. Our finding of divergent effects of interneuron function in PFC is in line with previous studies showing opposing effects on emotionality in acute vs chronic somatostatin (SST) IN inhibition in the PFC (Soumier and Sibille, 2014) and on auditory information processing in acute vs chronic IN inhibition in auditory cortex (Seybold et al., 2012). Our data suggests distinct neuronal ensembles and brain circuitry may be involved in modulating acute vs chronic stress mediated behavioral outcomes. It is also possible that chronic stress may result in plastic changes in the same neuronal ensemble recruited by acute stress, leading to differences in stress response. However, it is not known what specific plasticity in the neural network underlies the emergence of opposing phenotypes following chronic stress and further studies are needed to investigate the mechanisms.

There are few caveats to the present study that must be considered in the interpretation of the data. In this study we did not observe alterations in stress coping strategies in FST in our CVS group compared to controls. This is due to high rates of immobility in our control animals. Typically immobility duration in control mice should be around 60% (Wohleb et al., 2016), but in our control mice immobility was around 73%. It is known that body weight and age of rodents has a significant effect on behavior in the FST (Bogdanova et al., 2013; Hryhorczuk et al., 2013). It is possible that the high body weight and age of our mice resulted in a predominantly floating behavior leading to a ceiling effect, reducing the window to detect an increase in immobility typically observed after CVS exposure. Our control No CVS rats also received chronic injections. Chronic injection stress increases glutamate levels in the brain (Moghaddam and Bolinao, 1994) and might be acting as a stressor which may explain the greater immobility behavior observed in our controls. Nevertheless, we do see CVS effects in body weight, organ weights and in reduced Fos activation in stress regulatory brain regions demonstrating physiological effects of CVS in our study. This study was conducted only in male mice, to further prior research that showed alteration in inhibitory synaptic drive in the IL of males (McKlveen et al., 2019, 2016). Because PV IN modulation may have sex specific effects (Shepard et al., 2016), it would be important to examine the effects of PV IN modulation during stress in females. Finally, in this study we euthanized animals after exposure to one behavioral paradigm FST in order to get anatomical Fos expression dataset to a novel stressor following CVS. Chronic stress can be characterized by cellular and behavioral changes spanning multiple interconnected neural network adaptations which were not explored in our current study. In order to get a clear representation of how PV IN modulation during stress is affecting emotionality, additional behaviors may be worth exploring in follow-up experiments.

Taken together, our data are consistent with a causal role of IL PV INs in initiating and coordinating coping strategies and physiological outcomes in response to stress. Chronic stress-mediated hypoactivity and aberrant behavioral responses may be mediated partly via plastic changes in PV IN function and may play a role of stress-related pathologies (e.g., depression and PTSD). Our data indicate that chemogenetic inhibition of PV INs during chronic stress, which reduces PV-initiated inhibition in the context of each individual stressor experience, may block or attenuate inhibition of glutamatergic neurons. In this case, maintenance of glutamatergic excitability is sufficient to attenuate some (but not all) behavioral and physiological consequences of chronic stress exposure, including decreased passive (immobility) and increased active coping behaviors (swimming) in the FST, preventing CVS effects on reduction in neuronal Fos activity and in preventing adrenal hypertrophy. Taken together, our findings suggest that reducing the activity of PV INs in the PFC during chronic stress may facilitate output of prefrontal neurons and could provide therapeutic benefits for stress related disorders.

In conclusion, this study provides support that PV INs play a role in chronic stress mediated coping behaviors and physiological phenotypes. Furthermore, the study adds to the current knowledge regarding possible mechanisms of prefrontal hypofrontality, and how PV INs may be involved in driving chronic stress related pathologies. The study also highlights opposing effects of acute and chronic PV IN inhibition indicating different underlying mechanisms involved in acute vs chronic stress paradigms. Overall, this study shows that reducing PV IN activity to promote prefrontal output may be an effective treatment strategy for stress related illnesses.

## EXTENDED DATA FIGURE LEGENDS

**Figure 4-1.**
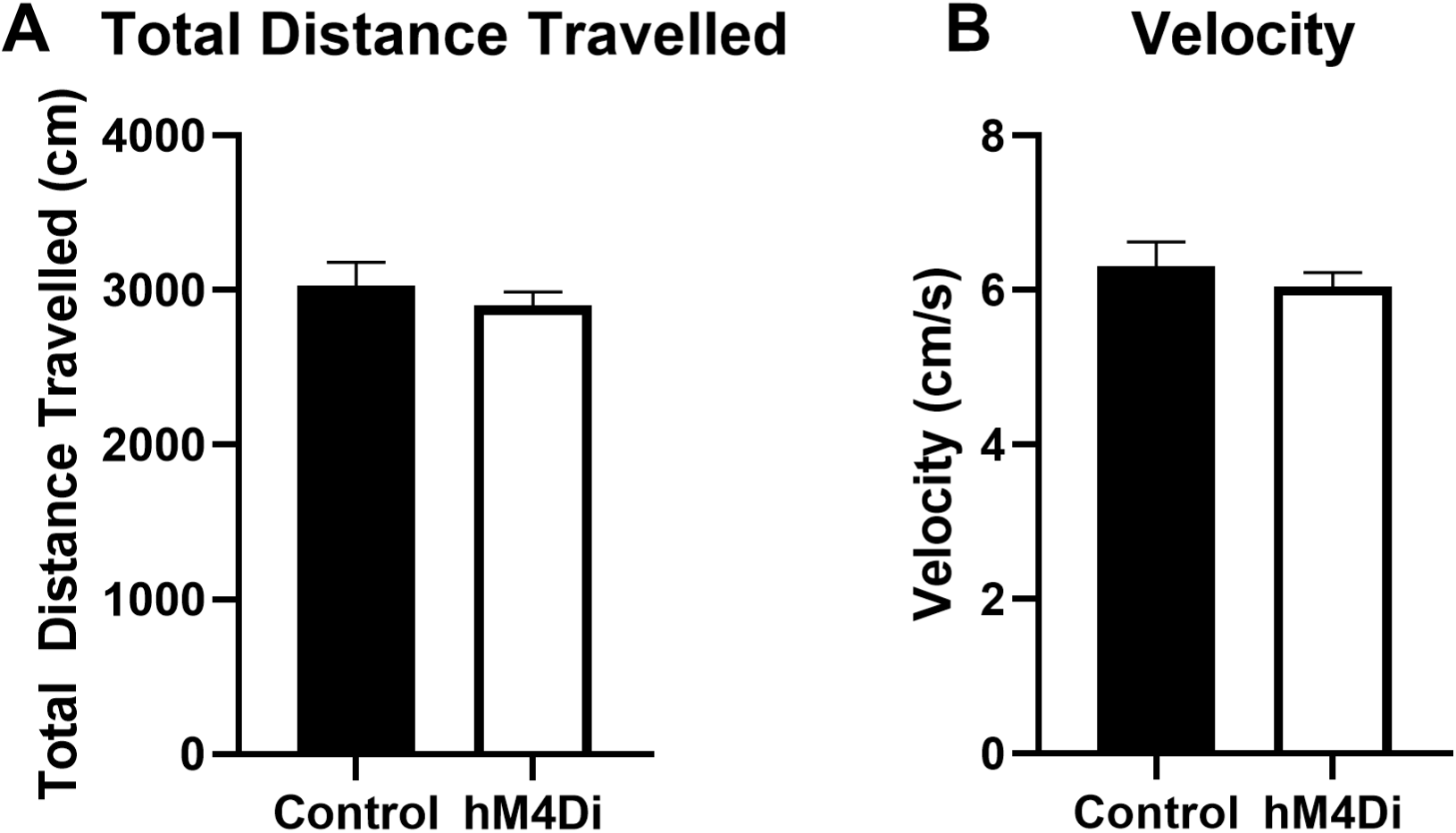
Effect of chronic inhibition of PV IN on Locomotor Activity in Y maze on day 12^th^ of CVS. Y maze was used as one of the stressors of the CVS paradigm. Chronic inhibition of PV INs had no effect on locomotor activity as demonstrated by no change in distance travelled (A) or velocity (B) in hM4Di group compared with control group. Values represent mean ± SEM, n = 9-10 per group (p>0.05).

**Figure 4-2.**
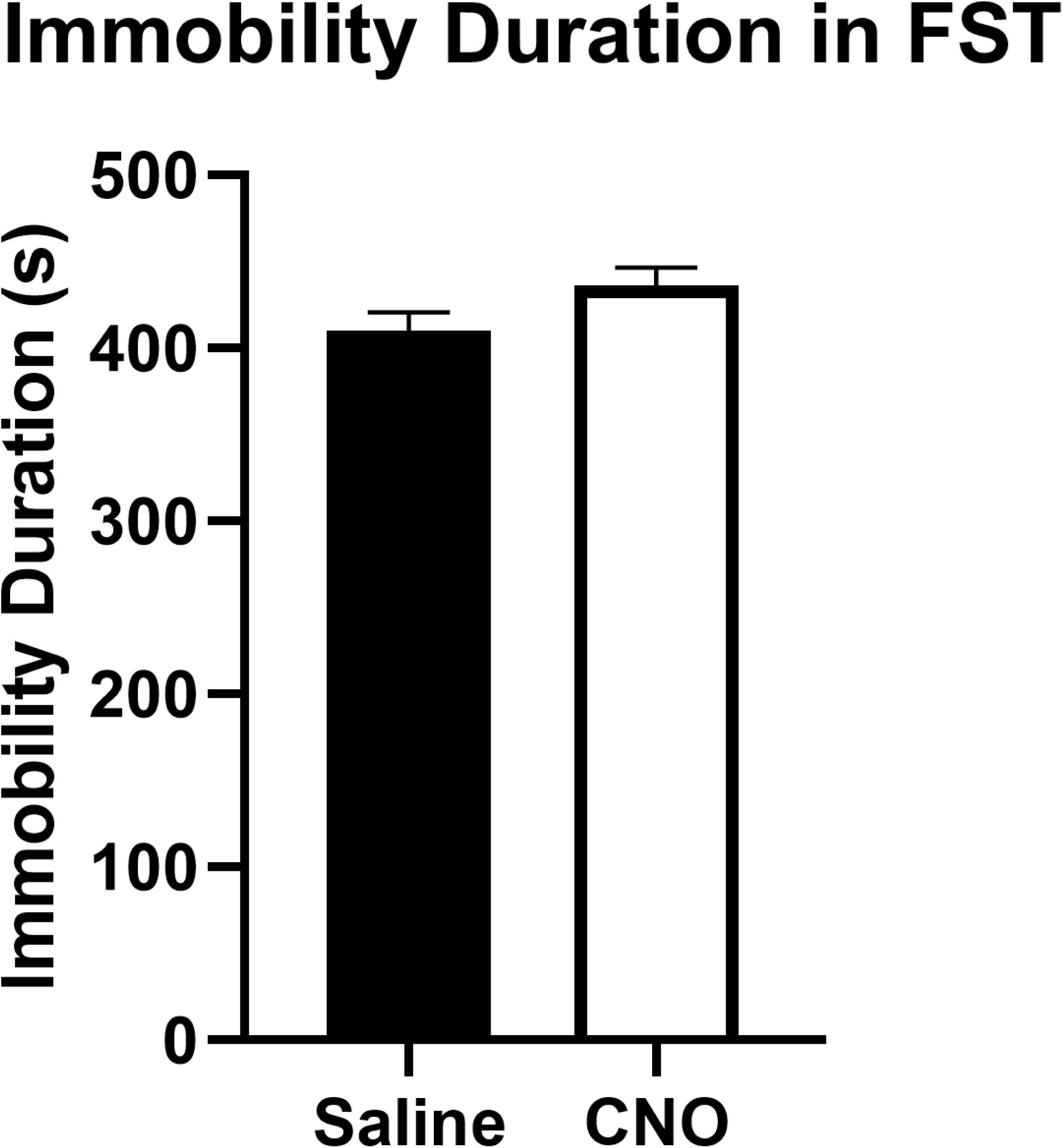
Effect of chronic dosing of CNO in control stress naive animals compared with saline in FST. Chronic CNO administration has no effect on immobility duration in forced swim test. Values represent mean ± SEM, n = 4 per group (p>0.05).

## EXTENDED DATA TABLE

**Table 5-1.** Expression of Fos protein in different brain regions following FST.

| Brain Regions | Groups | Fos Counts | F test |
| --- | --- | --- | --- |
| Lateral Septum | No CVS Control<br>CVS Control<br>No CVS hM4Di<br>CVS hM4Di | 122.2 +/- 6.8<br>113.1 +/- 4.1<br>127.1 +/- 9.2<br>108.1 +/- 7.7 | Main effect of stress only:<br>F (1,12)=7.2. p=0.02 |
| Anterior Bed Nucleus of Stria Terminalis (BNST) | NO CVS Control<br>CVS Control<br>No CVS hM4Di<br>CVS hM4Di | 76.8 +/- 5.0<br>66.7 +/- 8.1<br>73.0 +/-2.3<br>65.6 +/- 6.6 | No Effect of stress or treatment<br>Stress: F ( 1,14)=1.6; p=0.2<br>Treatment: F (1,14)=0.1; p=0.7 |
| Posterior Ventral BNST | NO CVS Control<br>CVS Control<br>No CVS hM4Di<br>CVS hM4Di | 79 +/- 8.1<br>62.1 +/-4.0<br>60.1 +/- 2.9<br>41.9 +/-4.1 | Main effect of Stress only:<br>F( 1,24)=12.4;p=0.002 |
| Dorsolateral PAG | NO CVS Control<br>CVS Control<br>No CVS hM4Di<br>CVS hM4Di | 51.7 +/- 3.7<br>52.6 +/-4.0<br>64.5 +/- 8.4<br>46.8 +/- 5.2 | No effect of Stress or Treatment<br>Stress: F ( 1,15)=2.1; p=0.2<br>Treatment: F (1,15)=0.4; p=0.6 |

Table depicts Fos protein expression in lateral septum, anterior and posterior ventral bed nucleus of stria terminalis (BNST) and dorsolateral periaqueductal gray (PAG). No significant treatment effects of PV IN inhibition was observed in any of the above mentioned brain regions. Values represent mean ± SEM, n = 9–10 per group.

